# Hemolysin liberates bacterial outer membrane vesicles for cytosolic lipopolysaccharide sensing

**DOI:** 10.1101/290445

**Authors:** Shouwen Chen, Dahai Yang, Ying Wen, Zhiwei Jiang, Lingzhi Zhang, Jiatiao Jiang, Yaozhen Chen, Tianjian Hu, Qiyao Wang, Yuanxing Zhang, Qin Liu

**Affiliations:** State Key Laboratory of Bioreactor Engineering, East China University of Science and Technology, Shanghai 200237, China; Laboratory for Marine Biology and Biotechnology, Qingdao National Laboratory for Marine Science and Technology, Qingdao 266071, China; Shanghai Engineering Research Center of Maricultured Animal Vaccines, Shanghai 200237, China; Department of Pathology and Comprehensive Cancer Center, University of Michigan, Ann Arbor, Michigan 48109, USA; Department of Transfusion Medicine, Xijing hospital, Xi’an 710032, China

**Keywords:** hemolysin, OMVs, noncanonical inflammasome, intestinal infection

## Abstract

Inflammatory caspase-11/4/5 recognize cytosolic LPS from invading Gram-negative bacteria and induce pyroptosis and cytokine release, forming rapid innate antibacterial defenses. Since extracellular or vacuole-constrained bacteria are thought to rarely access the cytoplasm, how their LPS are exposed to the cytosolic sensors is a critical event for pathogen recognition. Hemolysin is a pore-forming bacterial toxin, which was generally accepted to rupture cell membrane, leading to cell lysis. Whether and how hemolysin participates in non-canonical inflammasome signaling remains uncovered. Here, we show that hemolysin-overexpressed enterobacteria triggered significantly increased caspase-4 activation in human intestinal epithelial cells (IECs). Hemolysin promoted LPS cytosolic delivery from extracellular bacteria through dynamin-dependent endocytosis. Further, we revealed that hemolysin was largely associated with bacterial outer membrane vesicles (OMVs) and induced rupture of OMV-containing vacuoles, subsequently increasing LPS exposure to the cytosolic sensor. Accordingly, overexpression of hemolysin promoted caspase-11 dependent IL-18 secretion, gut inflammation, and enterocyte pyroptosis in orally-infected mice, which was associated with restricting bacterial colonization in vivo. Together, our work reveals a concept that hemolysin promotes noncanonical inflammasome activation via liberating OMVs for cytosolic LPS sensing, which offers insights into innate immune surveillance of dysregulated hemolysin via caspase-11/4 in intestinal antibacterial defenses.

**Significance:** Sensing of lipopolysaccharide (LPS) in the cytosol triggers non-canonical inflammasome-mediated innate responses. Recent work revealed that bacterial outer membrane vesicles (OMVs) enables LPS to access the cytosol for extracellular bacteria. However, since intracellular OMVs are generally constrained in endosomes, how OMV-derived LPS gain access to the cytosol remains unknown. Here, we reported that hemolysin largely bound with OMVs and entered cells through dynamin-dependent endocytosis. Intracellular hemolysin significantly impaired OMVs-constrained vacuole integrity and increased OMV-derived LPS exposure to the cytosolic sensor, which promoted non-canonical inflammasome activation and restricted bacterial gut infections. This work reveals the role of hemolysin in promoting non-canonical inflammasome activation and alerting host immune recognition, which provides insights into the more sophisticated biological functions of hemolysin upon infection.

## Introduction

The host innate immune system can sense invading bacteria by detecting pathogen-associated molecular patterns (PAMPs) (Vance *et al*, 2009). Lipopolysaccharide (LPS), a component of the outer cell membrane of Gram-negative bacteria, is one of the strongest immune activators (Rosadini & Kagan, 2017). Extracellular and endocytosed LPS is recognized by the transmembrane protein Toll-like receptor 4 (TLR4), leading to gene transcriptional regulation in response to infection (Kagan *et al*, 2008). Recent studies showed that host can detect LPS in the cytosol via a second LPS receptor, caspase-11 in mice and caspase-4/5 in humans (Hagar *et al*, 2013; Kayagaki *et al*, 2013; Shi *et al*, 2014). Caspase-11/4/5 directly binds cytosolic LPS (Shi *et al*, 2014), leading to its own activation, which thus cleaves gasdermin D to induce pyroptotic cell death and activate non-canonical activation of NLRP3 to release interleukin-1β (IL-1β) or IL-18 (Shi *et al*, 2015; Yang *et al*, 2015). Therefore, compartmentalization of LPS receptors within cells allows host to respond differentially and sequentially to LPS at distinct subcellular locales, which function in concert to constitute host noncanonical inflammasome defenses.

Caspase-11/4/5, as cytosolic sensors, only recognize LPS that has entered the host cell cytoplasm; however, the mechanism by which LPS from invading bacteria gains access to the cytosolic sensors remains unclear. For intracellular bacteria, although some bacteria such as *Burkholderia* (Aachoui *et al*, 2013) are cytoplasm-residing and easily expose LPS to the cytosolic sensors, many other bacteria such as *Salmonella typhimurium* (Broz *et al*, 2012) or *Legionella pneumophila* (Case *et al*, 2013) are predominantly constrained in pathogen-containing vacuoles (PCVs), probably masking LPS from cytosolic innate sensing. In addition to vacuolar bacteria, many extracellular bacteria, including *Escherichia coli* (Kayagaki *et al*, 2011), *Vibrio cholerae* (Kailasan *et al*, 2014), *Citrobacter rodentium* (Meunier *et al*, 2014), and *Haemophilus influenzae* (Rathinam & Fitzgerald, 2016) are thought to rarely access the cytoplasm, but induce caspase-11 dependent pyroptosis and cytokine release in cells. Thus, LPS entry into cell cytoplasm is a critical event for recognition of vacuolar or extracellular bacteria by non-canonical inflammasome. First, vacuolar bacteria may shed their LPS from endosome into the cytosol (Garcia-del *et al*, 1997). Lipopolysaccharide-binding protein (LBP) is also implicated in facilitating intracellular LPS delivery (Kopp *et al*, 2016). Alternatively, guanylate-binding protein 2 (GBP2) induces lysis of PCVs and promotes LPS leakage into the cytoplasm (Finethy *et al*, 2015; Meunier *et al*, 2014; Pilla *et al*, 2014). Recently, Vijay A.K. Rathinam and his colleagues improved the understanding of how non-canonical inflammasomes detect extracellular Gram-negative bacteria. Briefly, the outer membrane vesicles (OMVs) of extracellular bacteria enter cells by dynamin-dependent endocytosis, enabling LPS to access the cytosol by escaping from early endosomes (Vanaja *et al*, 2016). However, how OMVs gain access from early endosomes to the cytosol remains unknown.

Hemolysin belongs to the pore-forming protein family, rupturing the cell membrane and leading to cell lysis at high doses (Wiles & Mulvey, 2013). Recently, hemolysin was found to participate in modulating cell death pathways at sublytic concentrations (Bielaszewska *et al*, 2013; Ristow & Welch, 2016; Wiles & Mulvey, 2013), representing more sophisticated toxin activity in contrast to outright pore-forming function. Uropathogenic *Escherichia coli* (UPEC) isolate CP9 activates caspase-3/7 and stimulates rapid cell apoptotic death in vitro; this phenotype was lost in a Δ*hlyA* mutant (Russo *et al*, 2005), indicating the involvement of hemolysin in the cell apoptosis pathway. Recently, increasing evidence suggests that hemolysin promotes activation of inflammasome signals during infection. For example, entero-hemolysin of enterohemorrhagic *E. coli* (EHEC) O157:H7 triggered mature IL-1β secretion in human macrophages (Zhang *et al*, 2012), and α-hemolysin of UPEC CFT073 mediated NLRP3-dependent IL-1β secretion in mouse macrophages (Schaale *et al*, 2016). Strikingly, overexpression of hemolysin in UPEC UT189 activated significantly increased caspase-4 dependent cell death and IL-1α release than the controls (Nagamatsu *et al*, 2015). This is the first evidence demonstrating the relevance of hemolysin in caspase-4 activation, indicating that hemolysin might contribute to non-canonical inflammasome activation.

In this study, we first demonstrated that hemolysin in various enterobacteria significantly promoted caspase-4 dependent pyroptosis and IL-18 secretion in IECs. Further, we provided insights into the mechanism of hemolysin-mediated increase in the sensitivity of non-canonical inflammasome to invading bacteria. We showed that hemolysin internalizes into cells via binding to OMVs and promotes rupture of OMV-containing vesicles, thereby releasing OMV-derived LPS into the cytoplasm and eventually triggering significant activation of non-canonical inflammasomes in IECs. Oral infection of mice showed that abnormal expression of hemolysin in vivo alerts the immune system and induces caspase-11-dependent enterocyte pyroptosis and IL-18 secretion, which significantly constrains bacterial infection in the gut. Collectively, our results reveal that hemolysin enables OMV-mediated LPS cytosolic delivery for caspase-11/4 sensing, which alarms intestinal innate immune surveillance in vivo, providing insights into the manipulation of non-canonical inflammasome signals by invading bacteria.

## Results

### Hemolysin promotes caspase-4 dependent cell death and IL-18 secretion during infection

To screen for bacterial factors involved in regulating non-canonical inflammasome activation, a gene-defined mutant library of *Edwardsallar tarda* (*E. tarda*), an enteric pathogen infecting hosts from fish to human (Chen *et al*, 2017; Wang *et al*, 2009), was used to identify mutants that induced significantly increased pyroprosis in HeLa cells. Compared to the wild-type strain (EIB202), one of the mutants (0909I) greatly increased LDH release (Fig. S1A and S1B) and IL-18 secretion (Fig. S1C), accompanied by significantly induced caspase-4 activity (Fig. S1D) in HeLa. To explore whether 0909I promotes caspase-4-dependent non-canonical inflammasome activation in IECs, we extended the bacterial infection experiments to the wild-type and *Caspase-4*^−/−^ Caco-2 and HT-29 cells. Robust proptosis (Fig. 1A and Fig. S1E) and significantly increased LDH release (Fig. 1B and Fig. S1F) and IL-18 secretion (Fig. 1C and Fig. S1G) were detected in wild-type cells infected with 0909I compared to those infected with EIB202, which were counteracted in *Caspase4*^−/−^ cells. These data indicate that *E. tarda* mutant 0909I promotes caspase-4 dependent inflammasome activation in non-phagocyte cells.

**Figure 1.**
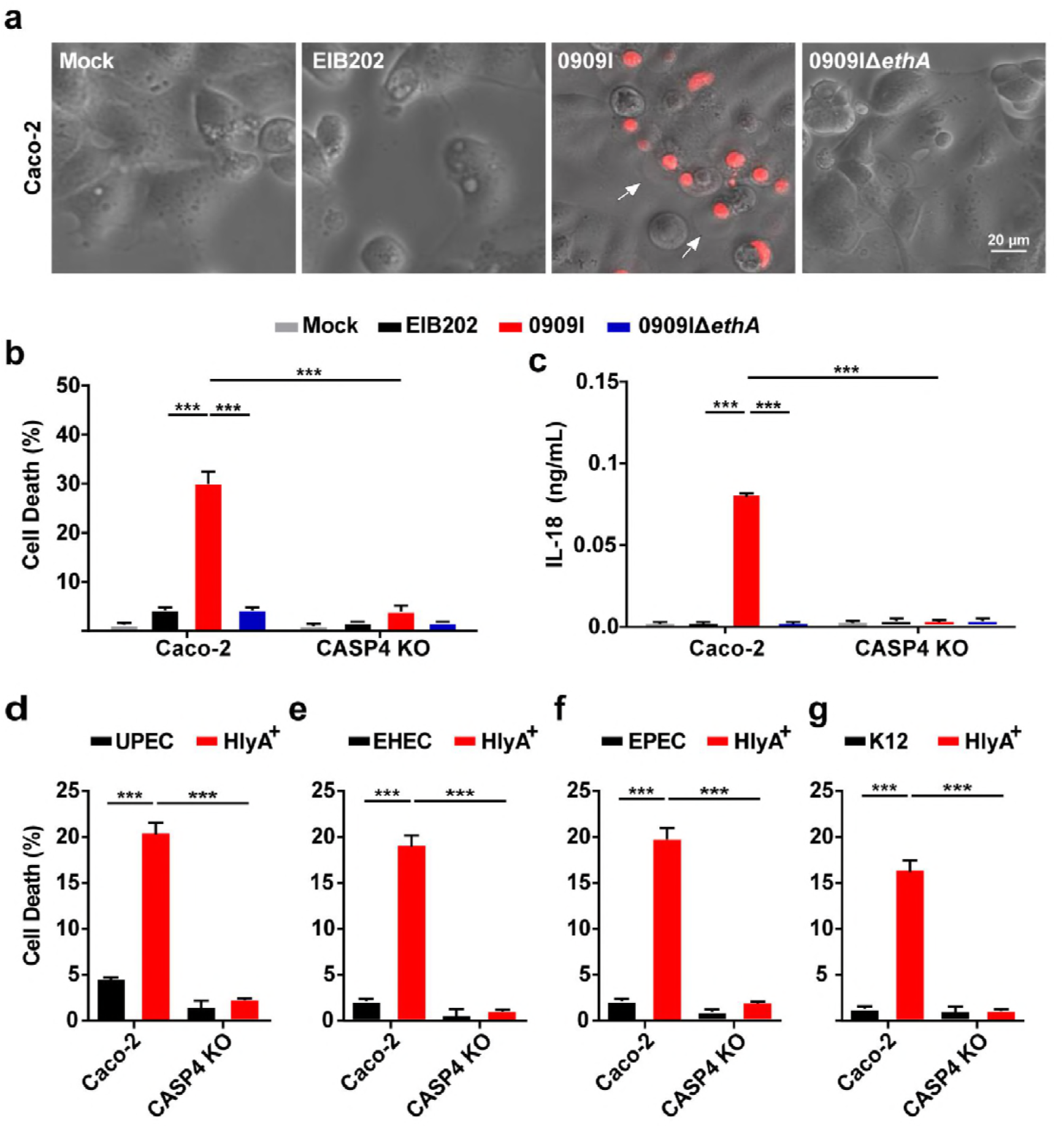
Hemolysin-overexpressed strains promotes caspase-4 activation during infection. (A) Cell morphology with propidium iodide (PI) staining of wild-type Caco-2 cells infected with the indicated *E. tarda* strains (MOI = 25, 4 hpi). Scale bar, 20 μm; arrows, pyroptotic cells. (D and F) LDH release or IL-18 secretion detected in wild-type and *Caspase-4*^−/−^ Caco-2 cells infected with the indicated *E. tarda* (MOI = 25, 4 hpi) (B and C) or *E. coli* strains (MOI = 50, 4 hpi) (D-G). Graphs show the mean and s.e.m. of triplicate wells and are representative of three independent experiments. *\*p* < 0.05, *\*\*p* < 0.01, *\*\*\*p* < 0.001; NS, not significant (two-tailed t-test).

Next, bioinformatics analysis revealed that the transposon insert site within 0909I is located upstream of a non-RTX hemolysin-encoding gene, *ethA* (Wang *et al*, 2010). In agreement with the upregulated transcription level of *ethA* (Fig. S2A), 0909I showed higher EthA expression (Fig. S2B) and bacterial hemolytic activity (Fig. S2C) than EIB202, which were abolished in the strain of 0909IΔ*ethA*. Further, deletion of *ethA* significantly impaired the ability of 0909I to increase caspase-4 activation in Caco-2 (Fig. 1A-1C) and HT-29 cells (Fig. S1E-S1G). These data suggest that 0909I promotes non-canonical inflammasome activation by increasing hemolysin expression.

To explore whether hemolysin also upregulates non-canonical inflammasome activation in other enteric bacteria, we expanded the investigation to the best-known RTX hemolysin, HlyA in UPEC, enterohemorrhagic *E. coli* (EHEC), enteropathogenic *E. coli* (EPEC) and *E. coli* K12. Similarly, the *E. coli* strains, containing HlyA expression plasmid, showed significantly higher hemolytic activity (Fig. S2D), which elicited a higher level of caspase-4 dependent LDH release (Fig. 1D-1G) and IL-18 maturation (Fig. S3A-3D) than their corresponding wild-type strains. Together, these results suggest that hemolysin plays critical roles in promoting caspase-4 activation during Gram-negative bacterial infection.

### Hemolysin promotes LPS cytoplasmic release through dynamin-dependent endocytosis

Host immunity senses bacterial LPS in the cytosol via caspase-11/4/5 (Hagar *et al*, 2013; Kayagaki *et al*, 2013; Shi *et al*, 2014). Because extracellular LPS cannot simply diffuse across the membrane, and most Gram-negative bacteria are not cytosolic, delivery of LPS from bacteria to the cytoplasm is thus very critical for non-canonical inflammasome activation. Because our data demonstrated that hemolysin increased caspase-4 activation in IECs, it is reasonable to speculate that hemolysin may promote cytosolic release of bacterial LPS during infection. We extracted the cytosol of uninfected or infected Caco2 cells using digitonin and assessed LPS levels in a limulus amebocyte lysate (LAL) assay. Digitonin is commonly used to isolate cytosol (Ramsby & Makowski, 2011), and the use of an extremely low concentration of digitonin (0.005%) for a very short duration allows extraction of cytosol devoid of plasma membrane, early and late endosomes, and lysosomes (Vanaja *et al*, 2016). The LAL assay showed that LPS was present in the cytosol of infected cells, but not in uninfected cells (Fig. 2A). Significantly higher levels of cytosolic LPS was detected in 0909I-infected cells than in EIB202-infected cells, and it was greatly reduced by deletion of *ethA* in 0909I (Fig. 2A). These data demonstrate that hemolysin promotes LPS delivery into the cytoplasm upon *E. tarda* infection.

**Figure 2.**
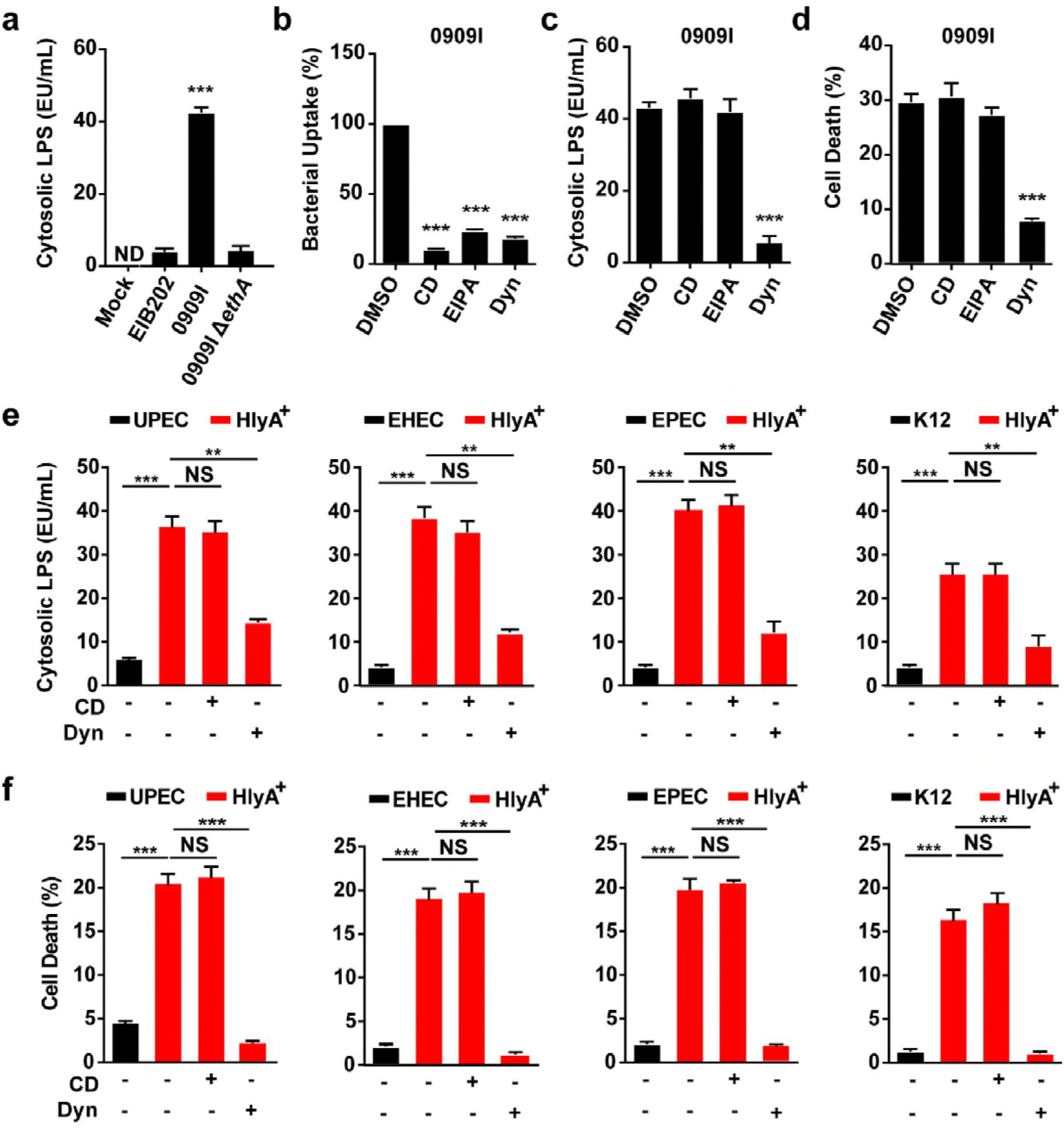
Hemolysin promotes LPS cytosolic release through Dynamin-dependent endocytosis. (A, C and E) LPS quantity by LAL assay in the cytosol extracted by digitonin fractionation from *Caspase-4*^−/−^ Caco-2 cells incubated with indicated *E. tarda* strains (MOI = 25, 4 hpi) (A, C) or *E. coli* strains (MOI = 50, 4 hpi) (E) in the presence of CD (10 μM), EIPA (30 μM), Dyn (80 μM), or not. (B) Bacterial count by agar plating in the cell pellets from *Caspase-4*^−/−^ Caco-2 cells incubated with indicated *E. tarda* 0909I (MOI = 25, 4 hpi), following treatment with 300 μg/mL gentamicin for 1 h to kill extracellular bacteria. (D, F) LDH release detected in wild-type Caco-2 infected with indicated *E. tarda* (MOI = 25, 4 hpi) (D) or *E. coli* strains (MOI = 50, 4 hpi) (F). Graphs show the mean and s.e.m. of triplicate wells and are representative of three independent experiments. *\*p* < 0.05, *\*\*p* < 0.01, *\*\*\*p* < 0.001; NS, not significant (two-tailed t-test).

*Edwardsallar tarda* is an intracellular pathogen, that replicates within a *E. tarda*-containing vacuole (ECV) (Cao *et al*, 2018; Hou *et al*, 2017; Vanaja *et al*, 2016), and has no direct access to the cytoplasm. We explored how hemolysin promotes LPS delivery from this bacterium to the cytosol. First, *E. tarda* strains showed similar internalization capacities between 0909I, EIB202, and 0909IΔ*ethA* (Fig. S4A), indicating that hemolysin did not affect cellular uptake of *E. tarda*. GBP2-induced vacuole destabilization was reported to release vacuolar bacteria for cytosolic LPS recognition (Meunier *et al*, 2014). However, no bacteria were recovered by agar plating in the cytosol extracted from *E. tarda*-infected cells, while *legionella pneumophila* Δ*sdhA* was used as a control of vacuole-escaping bacteria (Fig. S4B), suggesting that hemolysin did not promote vacuole-constrained *E. tarda* to enter the cytosol. Further, to discriminate that hemolysin promotes LPS release to the cytoplasm from vacuolar or extracellular bacteria, three endocytosis inhibitors, cytochalasin D (CD), 5-(N-ethhyl-n-isopropil)-amiloride (EIPA), and dynasore (Dyn) were used to inhibit the internalization of *E. tarda* in Caco-2 cells. Clearly, CD and EIPA inhibited the internalization of 0909I (Fig. 2B), but did not reduce either cytoplasmic LPS release (Fig. 2C) or caspase-4 activation (Fig. 2D and Fig. S5A), suggesting that hemolysin-mediated LPS delivery predominantly depends on extracellular bacteria. Unexpectedly, dynasore similarly inhibited cellular uptake of *E. tarda* (Fig. 2B), but significantly suppressed LPS delivery (Fig. 2C) and sensing by cytosolic caspase-4 (Fig. 2D and Fig. S5A), indicating that dynasore may block a pathway that delivers LPS from extracellular bacteria to the cytoplasm. Next, we investigated the effects of CD and Dyn in *E. coli*-infected Caco-2 cells. Dyn significantly repressed LPS cytosolic release (Fig. 2E), cell death (Fig. 2F) and IL-18 secretion (Fig. S5B) in cells, while CD showed no influence on them. These results suggest that hemolysin promotes LPS cytosolic release from extracellular bacteria through a dynamin-dependent endocytosis process.

### Hemolysin-mediated caspase-4 activation is dependent on association with OMVs

Outer membrane vesicles (OMVs) are spherical, bilayered nanostructures constitutively released by growing bacteria (Schwechheimer & Kuehn, 2015). The association of bacterial toxins with OMVs protects them from inactivation or degradation during infection, representing a highly efficient mechanism of bacteria modulating host defenses (Kaparakis-Liaskos & Ferrero, 2015). Bacterial hemolysins, such as HlyA in *E. coli* largely bind OMVs and enter cells by dynamin-dependent endocytosis (O’Donoghue & Krachler, 2016). To dissect the role of *E. tarda* hemolysin in promoting caspase-4 inflammasome activation, we first explored the localization of EthA in this bacterium. Here, blebbing and shedding of OMVs were observed in growing *E. tarda* (Fig. S6A) and produced in supernatants over time (Fig. S6B). We fractionated the bacterial culture into pellets, OMV-free supernatants and OMVs. Notably, over 60% of the total EthA was detected in the fraction of OMVs (Fig. 3A), indicating that OMV-associated EthA is the major form of this toxin in *E. tarda*. In accordance with the increased hemolytic activity in 0909I (Fig. S2C), significantly higher levels of EthA and hemolytic activity were detected in 0909I OMVs than in EIB202 or 0909IΔ*ethA* OMVs (Fig. 3B). In contrast, when subject to proteinase K (PK) digestion, in which proteins inside OMVs are protected from degradation, OMV-associated EthA was completely degraded, which agrees with the decreased hemolytic activity (Fig. 3B), suggesting that EthA is exposed on the exterior of *E. tarda* OMVs.

**Figure 3.**
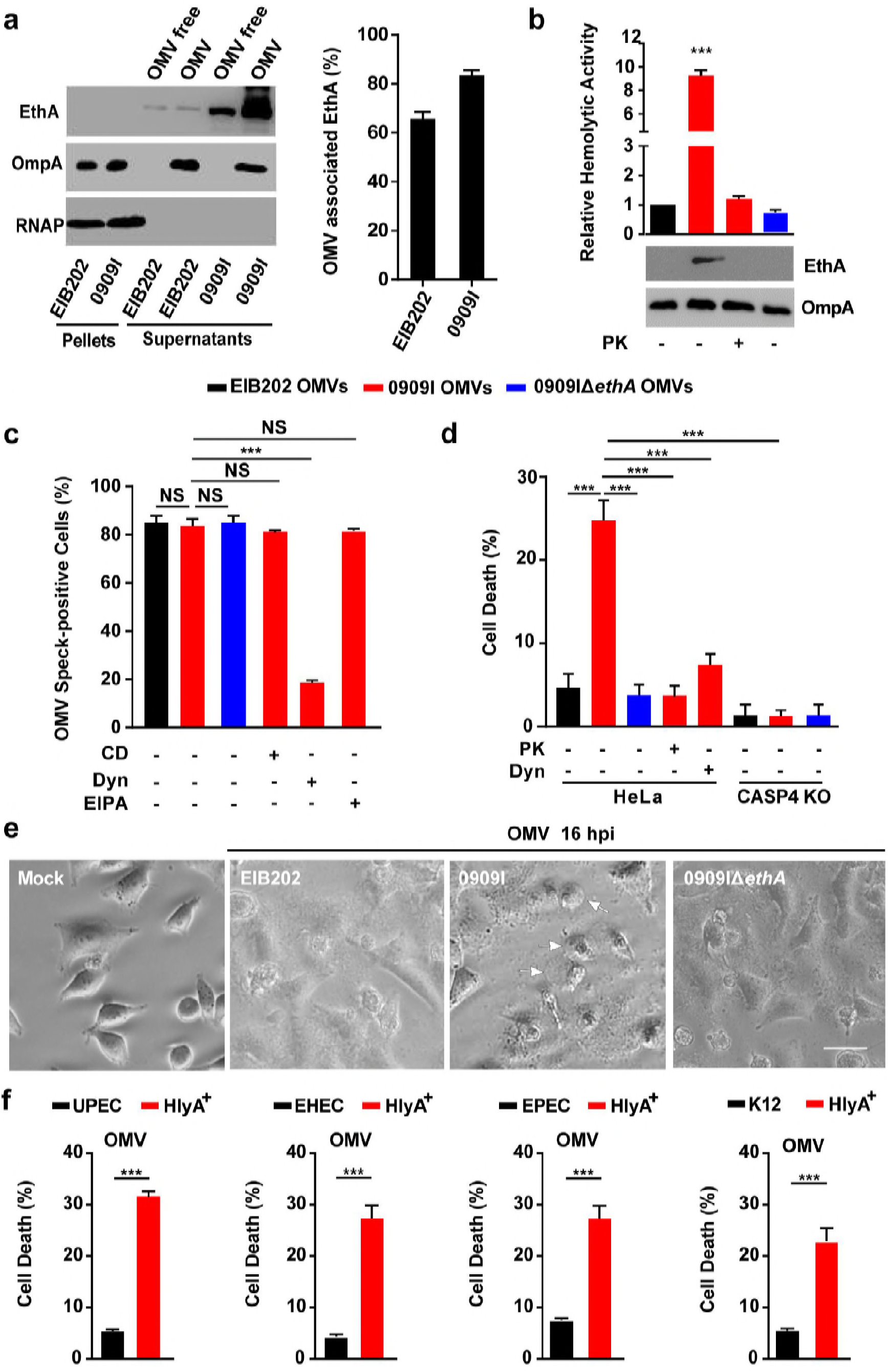
Hemolysin associates with OMVs to promote caspase-4 activation. (A and B) Immunoblots for EthA in different fractions of *E. tarda* culture at 10 h post-inoculation (A) and assay for hemolytic activity and immunoblots for EthA in OMVs, treated with PK (5 μg/mL, 5 min) or not (B). (C) Immunostaining for intracellular OMVs using anti-OmpA antibody and quantification of OMV speck-positive cells in HeLa cells incubated with indicated *E. tarda* OMVs (20 μg/1 × 10^5^ cells, 4 h) in the presence of CD (10 μM), EIPA (30 μM), Dyn (80 μM), or not. Percentage of cells containing OMV specks was calculated for at least 500 cells. (E) Morphology of HeLa cells infected with indicated *E. tarda* OMVs (50 μg/1 × 10^5^ cells, 16 h). Scale bar, 20 μm; arrows, pyroptotic cells. (D and F) LDH release detected in wild-type or *Caspase-4*^−/−^ HeLa cells incubated with purified *E. tarda* OMVs (50 μg/1 × 10^5^ cells, 20 h) pretreated with PK (5 μg/mL, 5 min), Dyn (80 μM), or not (D), or purified *E. coli* OMVs (50 μg/1 × 10^5^ cells, 20 h) (F). Graphs show the mean and s.e.m. of triplicate wells and are representative of three independent experiments. *\*p* < 0.05, *\*\*p* < 0.01, *\*\*\*p* < 0.001; NS, not significant (two-tailed t-test).

A recent study suggested that OMVs were responsible for delivering LPS into the cytosol and activating caspase-11 inflammasome (Vanaja *et al*, 2016). Because *E. tarda* hemolysin was largely associated with OMVs and critically involved in caspase-4-dependent inflammasome activation, it raises the possibility that *E. tarda* employs hemolysin-associated OMVs to activate the non-canonical inflammasome during infection. Subsequently, purified *E. tarda* OMVs were incubated with wild-type or *Caspase-4*^−/−^ cells. OMVs internalized and formed obvious specks within cells (Fig. S6C) at comparable level between EIB202, 0909I, and 0909IΔ*ethA*, which were significantly inhibited by treatment with Dyn, but not with CD or EIPA (Fig. 3C). Importantly, according to the higher hemolytic activity (Fig. 3B), 0909I OMVs induced obvious cell pyroptosis (Fig. 3D), and a higher level of LDH release (Fig. 3F) than EIB202 OMVs in wild-type cells, and this difference was counteracted in *Caspase-4^−/−^* cells (Fig. 3E). Further, hemolysin depletion by deleting *ethA* or pretreating OMVs with PK remarkably reduced 0909I OMV-induced caspase-4 activation (Fig. 3E). Simultaneously, treatment with dynasore to inhibit cellular uptake of 0909I OMVs reduced LDH release in wild-type cells (Fig. 3E). Similarly, purified *E. coli* HlyA^+^ OMVs significantly increased cell death than wild-type OMVs (Fig. 3F). These data suggest that hemolysin promotes OMV-mediated caspase-4 inflammasome activation in Gram-negative bacteria.

### Hemolysin impairs OMVs-containing vacuole integrity for cytosolic access of LPS

As LPS-enriched OMVs are internalized via endocytosis and restrained within endosomes (O’Donoghue & Krachler, 2016), it raises the question of how vacuole-contained OMVs achieve cytosolic localization of LPS. Because hemolysin is a type of bacterial pore-forming toxins, that leads to membrane permeabilization (Wiles & Mulvey, 2013), a possible explanation could be that hemolysin induces lysis of OMV-containing vacuoles (OCVs) and facilitates cytosolic exposure of LPS. Galectin-3 is a β-galactoside binding protein, that is specifically recruited to disrupted pathogen-containing vacuoles (Paz *et al*, 2010). We assessed the recruitment of GFP-tagged galectin-3 in cells incubated with OMVs. Indeed, 0909I OMVs induced more galectin-3 specks than EIB202 OMVs (Fig. S7A and Fig. 4A), and galectin-3 showed clearly association with OMV specks within wild-type cells (Fig. 4B). In contrast, removal of hemolysin from 0909I OMVs by proteinase K degradation or deleting *ethA* significantly decreased the intracellular galectin specks in wild-type cells (Fig. 4A). These data suggest that hemolysin promotes rupture of OMV-containing vacuoles.

**Figure 4.**
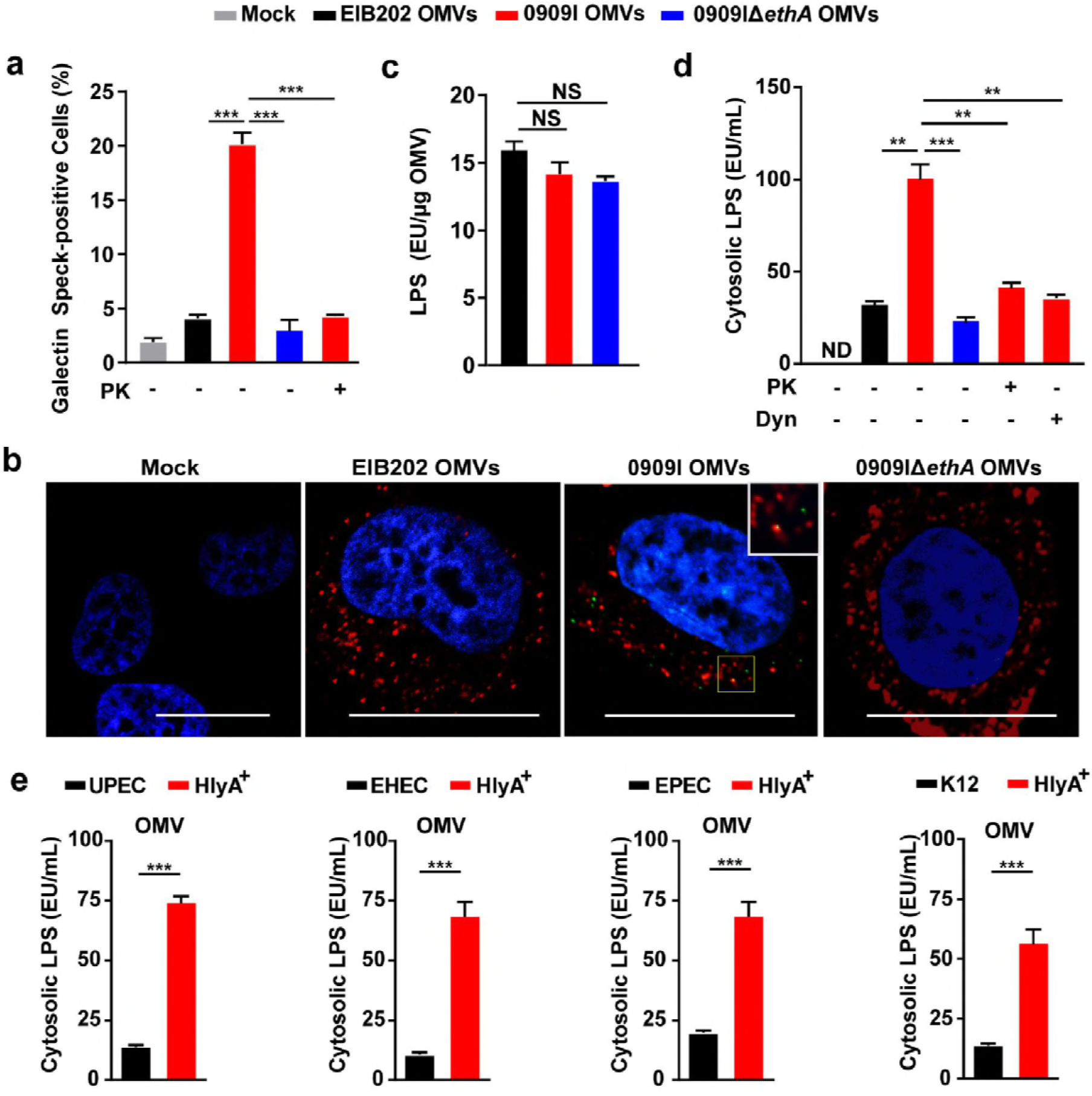
Hemolysin impairs OMVs-containing vacuole integrity for cytosolic access of LPS. (A) Quantification of galectin speck-positive cells in HeLa cells expressing GFP-tagged gelectin-3, incubated with *E. tarda* OMVs (50 μg/1 × 10^5^ cells, 20 h) treated by PK (5 μg/mL, 5 min) or not. Percentage of cells containing galectin specks was calculated for at least 500 cells. (B) Observation of intracellular galectin (green)-associated OMV specks using anti OmpA antibody (red). Scale bar, 20 μm; the white box indicates the co-localized signal between galectin and OMVs. (C) LPS quantification by LAL assay in purified E. tarda OMVs. (D and E) LPS quantification by LAL assay in the cytosol extracted by digitonin fractionation from *Caspase-4*^−/−^ HeLa cells, incubated with E. tarda OMVs (50 μg/1 × 10^5^ cells, 20 h) treated with PK (5 μg/mL, 5 min), Dyn (80 μM), or not (D), or *E. coli* OMVs (50 μg/1 × 10^5^ cells, 20 h) (E). Graphs show the mean and s.e.m. of triplicate wells and are representative of three independent experiments. *\*p* < 0.05, *\*\*p* < 0.01, *\*\*\*p* < 0.001; NS, not significant (two-tailed t-test).

Next, we explored the contribution of hemolysin to damaging the membrane of OCVs during *E. tarda* infection. As expected, 0909I triggered significantly increased signal of galectin aggregation in DMSO-treated cells, compared to EIB202 or 0909IΔ*ethA* (Fig. S7B and S7C). In the presence of EIPA, which inhibited the internalization of bacteria, but not OMVs into cells, 0909I induced obvious cytoplasmic galectin specks (Fig. S7B and S7C), indicating that 0909I-induced galectin aggregation is mainly associated with internalized OMVs rather than bacteria. Further, 0909I trigged a significant increase in cytoplasmic galectin signal, compared to EIB202 or 0909I Δ*ethA*, which was greatly suppressed by pretreating the cells with Dyn (Fig. S7B and S7C). These results indicate that hemolysin contributes to the destruction of OMV-residing vesicles during bacterial infection.

Because hemolysin triggered significant membrane rupture of OCVs within cells, we evaluated whether vesicle lysis facilitates LPS exposure to cytosolic sensors. We extracted the cytosol and quantified the cytosolic LPS in *Caspase-4^−/−^* cells upon incubation with purified OMVs. Although EIB202, 0909I, and 0909IΔ*ethA* OMVs showed comparable LPS contents (Fig. 4C) and uptake efficiencies (Fig. 3C and 3D), significantly increased cytosolic LPS was detected in the cytosol of 0909I OMV-incubated cells than in the controls. This difference was remarkably reduced by either eliminating OMV-bound hemolysin or damping OMV internalization (Fig. 4D). Furthermore, purified OMVs from *E. coli* strains were incubated with *Caspase-4^−/−^* cells for the cytosolic LPS assay. Accordingly, HlyA^+^ OMVs induced more LPS release into the cytoplasm than their correspondent wild-type OMVs (Fig. 4E), indicating that hemolysin promotes cytosolic release of OMV-LPS. Collectively, these data suggest that hemolysin-mediated membrane lysis of OCVs represents an important means to liberate OMV-LPS for cytosolic sensing in Gram-negative bacteria.

### Hemolysin increases recognition of invading bacteria by caspase-11 inflammasome in vivo

Intestinal epithelium cells are at the forefront of the host gut defense system. Accumulating data indicate the importance of IEC inflammasomes in shaping intestinal immune defense against bacterial infection via inducing IL-1α/β or IL-18 secretion or enterocyte pyroptosis (Sellin *et al*, 2015). As hemolysin was verified to promote non-canonical caspase-4 inflammasome activation in IECs, we further assessed the involvement of hemolysin in noncanonical inflammasome activation in vivo. C57BL/6 wild-type mice were orally infected with *E. tarda* strains. Compared to EIB202, 0909I showed significantly reduced bacterial burdens in the colon, cecum and lumen (Fig. 5A), but not at the systemic sites (Fig. S8A), indicating that that over-expressed hemolysin restricts *E. tarda* colonization in the mouse gut. Subsequently, it is interesting to explore the in vivo relevance of hemolysin-mediated gut infection restriction with non-canonical inflammasome activation. Caspase-11 is the murine ortholog of human caspase-4, which predominantly mediates IL-18 secretion and enterocyte pyroptosis (Knodler *et al*,2014; Pallett *et al*,2017) during intestinal infection. Notably, caspase-11 depletion counteracted the superiority of bacterial loads in wild-type mice infected with 0909I over EIB202 (Fig. 5A), suggesting the critical requirement of caspase-11 by hemolysin-triggered gut infection limitation.

**Figure 5.**
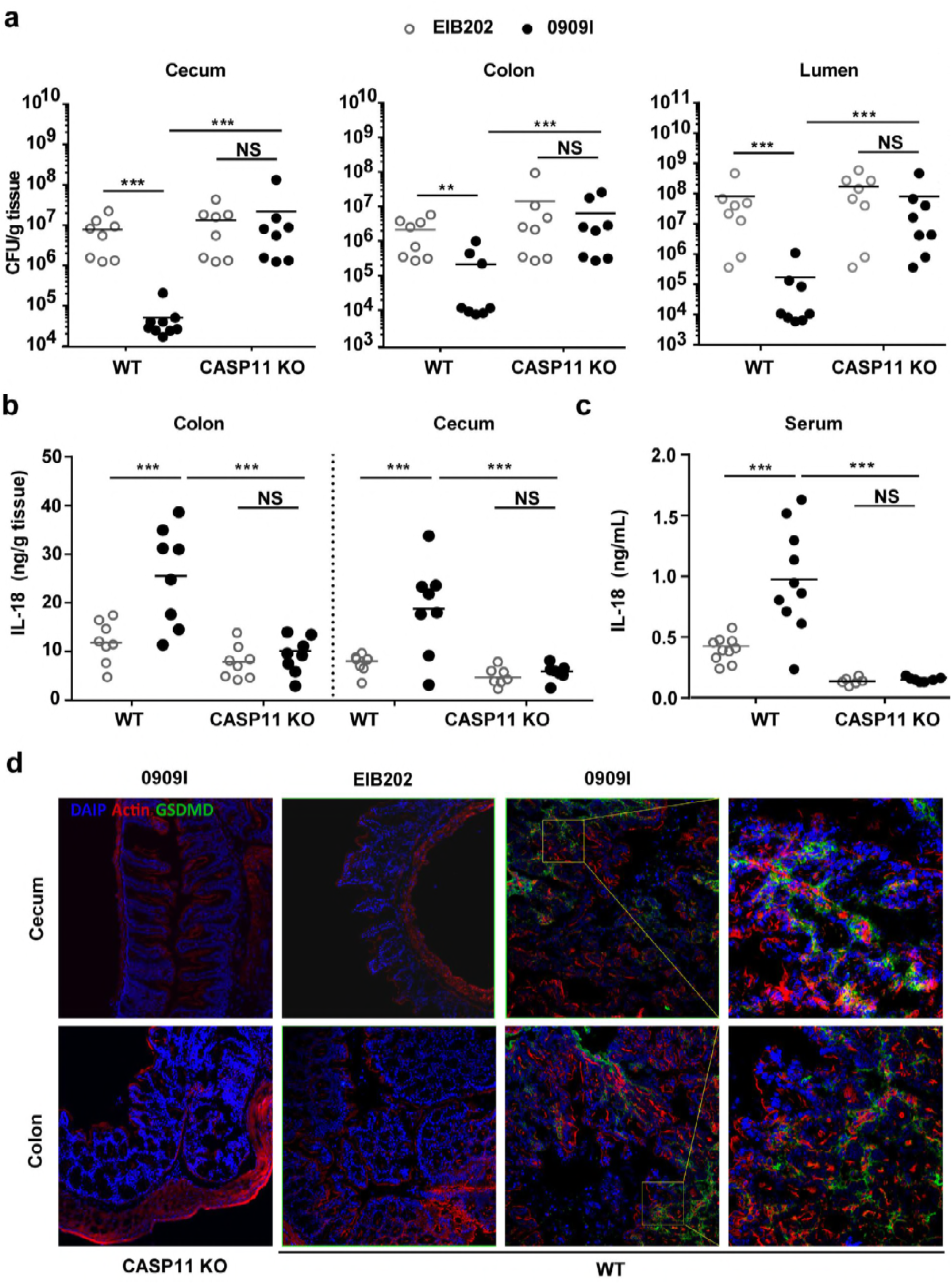
Hemolysin increases recognition of invading bacteria by caspase-11 in vivo. (A) Bacterial counting by agar plating in the colon, caecum, and lumen of wild-type or Caspase-11-/- mice orally-infected by EIB202 or 0909I (5 × 10^7^ cfu/g) at 24 hpi. (B and C) Quantification of IL-18 in the colon, cecum (B) and serum (C) of the mice described in a. (D) Confocal laser scanning of pyroptotic cells in the gut sections stained by anti-GSDMD antibody (green) without permeabilizing cells. The actin was stained in red and nuclei in blue; magnification = 20 ×; the boxes indicate the details of GSDMD signals in the guts. Graphs depict 6–10 mice per genotype and are representative of two to three independent experiments. *\*p* < 0.05, *\*\*p* < 0.01, *\*\*\*p* < 0.001; NS, not significant (one-way ANOVA).

Further, we demonstrated that 0909I induced remarkably higher mucosal and serum IL-18 level than EIB202 in wild-type mice (Fig. 5B and 5C). Tissue pathology analysis revealed that 0909I evoked prominent intestinal inflammation in wild-type mice, typically featured by focal filtration of inflammatory cells and epithelia cell shedding (Fig. S8B and S8C). In contrast, these phenotypes were largely neutralized in *Caspase-11*^−/−^ mice. Recent studies reported that inflammasome-induced pyroptosis caused enterocyte extrusion, which facilitates bacterial clearance (Knodler *et al*, 2014; Pallett *et al*, 2017; Sellin *et al*, 2015). To probe the pyroptotic cells in the intestinal epithelia, we immunoblotted for the cytosolic gasdermin D (GSDMD) in the gut sections without permeabilization. Obvious GSDMD-positive signals were observed in the colon and cecum of 0909I-infected wild-type mice, but absent in *Caspase-11*^−/−^ mice gut (Fig. 5D), indicating that 0909I induced membrane-rupturing pyroptosis in IECs and thus significantly increased enterocyte extrusion. Together, these results indicate that dysregulation of hemolysin in vivo alerts caspase-11 dependent intestinal defenses and restricts bacterial gut infection.

## Discussion

Sensing of LPS in the cytosol by inflammatory caspase-11/4/5 has emerged as a central event of innate immune responses during Gram-negative bacterial infections (Hagar *et al*, 2013; Kayagaki *et al*, 2013; Shi *et al*, 2015). Vanaja *et al*. reported that OMVs of extracellular Gram-negative bacteria can deliver LPS into the host cell cytosol from early endosomes; however, the mechanism of LPS translocation remains unclear. Biological membrane characteristics inherent to OMVs may permit them to fuse with endosomal membranes, leading to LPS cytosolic access (Vanaja *et al*, 2016). Recently, it was reported that GBPs were involved in OMV-dependent non-canonical inflammasome activation (Finethy et al, 2017; Santos et al, 2018). Mechanistically, GBPs didn’t promote the entry of OMVs into the cytosol, but directly target cytosolic OMVs and facilitate the interaction of LPS with caspase-11. In addition to host factors, whether bacterial factors, such as OMV-associated bacterial components, are involved in promoting non-canonical inflammasome activation remains unknown. Here, we demonstrate that hemolysin binds OMVs and promotes the lysis of OMV-residing vesicles, which facilitates cytosolic release of OMV-LPS and eventually triggers significant non-canonical inflammasome signals (Fig. 6). Our results suggest that hemolysin represents a biologically important mechanism for releasing endosome-constrained OMV-LPS to cytosolic sensors. In addition to hemolysin, whether other pore-forming proteins produced by bacteria (Bischofberger *et al*, 2012) play roles in liberating pathogen-associated molecular patterns for immune detection requires further examination.

**Figure 6.**
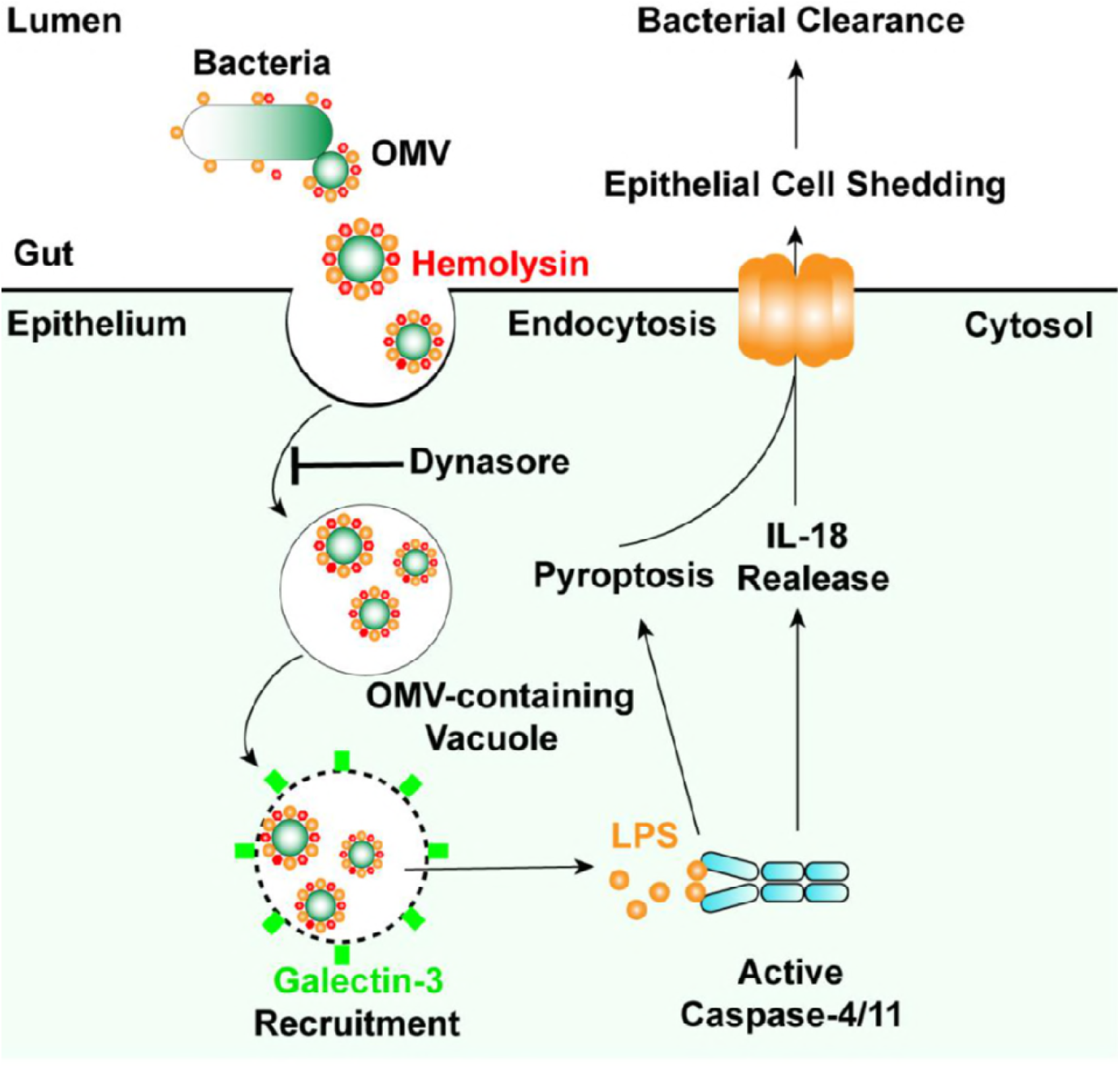
Model for bacterial hemolysin in regulating noncanonical inflammasome pathway. Hemolysin binds bacterial outer membrane vesicles (OMVs) and liberates OMV-associated LPS for caspase-4 sensing, which increases recognition of invading Gram-negative bacteria by non-canonical inflammasome and restricts bacterial gut infections.

Outer membrane vesicles are extruded from the surface of living bacteria and may entrap some of the underlying periplas (Schwechheimer & Kuehn, 2015). In addition to outer membrane proteins, LPS, phospholipids, and periplasmic constituents, OMVs deliver various bacterial cargos, including toxins intracellularly and further in specific compartments to modulate host defenses during infection (Kaparakis-Liaskos & Ferrero, 2015). Thus, OMVs offer bacterial products specific access to otherwise inaccessible cellular or tissue compartments. Particularly, OMVs enable the cytosolic localization of LPS for caspase-11 sensing (Vanaja *et al*, 2016), which is thought to be an important mechanism in extracellular bacteria. In this work, in addition to *E. coli* strains, the typical vacuolar bacterium, *E. tarda* (Zhang *et al*, 2016) also depends on OMVs to activate caspase-4 inactivation in IECs, indicating that OMV-mediated non-canonical inflammasome activation represents a more generalized mechanism for bacteria than previously thought. Moreover, because OMVs were linked to the translocation of diverse bacterial factors into host cells, it will be interesting to determine whether hemolysin also increases cytosolic recognition of other ligands such as DNA or flagellin by the AIM2 (Burckstummer *et al*, 2009) or NLRC4 (Zhao & Shao, 2015) inflammasome.

Hemolysin, as a class of pore-forming toxins, is known to disrupt the membrane of eukaryotic cells at high doses in vitro, which potentially facilitates bacterial invasion and dissemination during infection (Wiles & Mulvey, 2013). HlyA is the prototype member of the hemolysin family (Wiles & Mulvey, 2013). Only a very low percentage of nonpathogenic *E. coli* harbors HlyA, while ∼50% of UPEC strains harbor and express HlyA, presenting a close correlation between increased clinic *E. coli* pathogenesis with HlyA expression (Bielaszewska *et al*, 2014; Ristow & Welch, 2016; Wiles & Mulvey, 2013). In recent years, because it has been hypothesized that hemolysin is secreted at sublytic concentrations in vivo, there is increasing interest in understanding the more subtle effects of hemolysin on host cellular processes aside from outright lysis, including RhoA activation (Mansson *et al*, 2007), Akt signaling (Mansson *et al*, 2007), cell death pathway (Bielaszewska *et al*, 2013; Zhang *et al*, 2012) and protease degradation (Dhakal & Mulvey, 2012). Here, our study adds new findings to the scenario of hemolysin-mediated biological functions by identifying that abnormal expression of hemolysin in bacteria promotes host immune recognition of replicating pathogens via non-canonical inflammasomes and restricts bacterial colonization in vivo (Fig. 6). Because hemolytic proteins exhibit multilayered or even antagonistic roles in modulating host responses to infections, it is not surprising that bacteria have evolved complicated mechanisms to fine-tune the synthesis, maturation, and transport of hemolysin (Bielaszewska *et al*, 2014; Ristow & Welch, 2016; Wiles & Mulvey, 2013). Thus, studies of the spatiotemporal expression of hemolysin in vivo will be essential for understanding both host defense and bacterial survival.

Inflammasomes are activation platforms for inflammatory caspases, leading to pyroptosis and secretion of IL-1α, IL-1β, and IL-18 (Rathinam & Fitzgerald, 2016). The actions of inflammasomes have been well-studied in myeloid cells, and accumulating evidence highlights their role in non-professional immune cells acting as an important antibacterial defense mechanism (Sellin *et al*, 2015). Recent studies demonstrated that IECs utilize both canonical and non-canonical inflammasomes to promote mucosal defense against enteric bacterial pathogens (Knodler *et al*, 2014; Pallett *et al*, 2017; Sellin *et al*, 2014). These independent studies showed that pyroptosis leads to extrusion of infected enterocytes, which promotes bacterial infection in vivo even in the absence of IL-1 α, IL-1β, and IL-18. In our study, hemolysin overexpression significantly promoted caspase-11-dependent pyroptotic extrusion of enterocytes, correlating with decreased bacterial loads in vivo, supporting that cell intrinsic inflammasome-induced enterocyte extrusion is a generalized defense mechanism against acute bacterial infections. Further insights into whether enterocyte pyroptosis directly removes the replicative niches for pathogens or infectious agents released from pyroptotic cells leads to recruitment and priming of immune cells, ultimately contributing to pathogen clearance from the host, require future studies.

## Materials and Methods

### Cell culture and infections

Wild-type or *Caspase-4^−/−^* Caco-2, HT-29, and HeLa cells were incubated with indicated *E. tarda* or *E. coli* strains at a multiplicity of infection (MOI) of 25 or 50, respectively, or incubated with purified OMVs at 100 µg/10^5^ cells. Cell supernatants were harvested for IL-18 and LDH assays.

### Galectin assay

The plasmid expressing GFP tagged galectin-3 was constructed and introduced into HeLa cells by lentivirus transfection. Transfected cells were challenged with different *E*. *tarda* strains at a MOI of 50 or incubated with purified OMVs at 100 μg/10^5^ cells. Galectin speck formation within cells was observed under a confocal microscope.

### Cytosolic LPS quantification

*Caspase-4^−/−^* HeLa or Caco-2 cells post-incubation with the indicated strains or purified OMVs were treated with 0.005% digitonin. The extracted cytosol and residual fractions were subjected to the Limulus Amebocyte Lysate (LAL) assay to quantify LPS.

### Mice and Infections

C57BL/6J wild-type and *Caspase-11^−/−^* knockout mice (6-8 weeks old), pretreated with streptomycin, were orally infected with *E. tarda* strains at 5×10^7^ CFU/g. At 24 hours post-infection, indicated tissues or organs were collected and detected for bacterial loads, IL-18 secretion, gut histopathology, and enterocyte pyroptosis.

**A full description of methods used is provided in SI Methods**

## ACKNOWLEDGMENTS

We thank F. Shao for caspase-4 deficient HeLa cells and constructive suggestions, Y.F. Yao, D.P. Yan and M.A. Mulvey for providing *E. coli* strains. This work was supported by the National Natural Science Foundation of China No. 31622059 (Q.L.) and 31430090 (Y.Z.) and the Fundamental Research Funds for the Central Universities No. 222201717019 and 222201718004. Dahai Yang was supported by the Young Elite Scientists Sponsorship Program by CAST No. 2016QNRC001, Shanghai Chenguang Program No.16CG33 and Talent Program of School of Biotechnology in East China University of Science and Technology.

## Author contributions

Q.L. and S.C. designed research; S.C., D.Y., Y.W., Z.J., L.Z., T.H., J.J. and Y.C. performed research; Q.L., S.C., D.Y and Q.W. analyzed data; Q.L. and Y.Z. supervised research; and Q.L., S.C. and D.Y wrote the paper.

## Supplementary Information

### Supplementary Materials and Methods

#### Bacterial strains and plasmids

0909I *E. tarda* was screened from *E. tarda* gene-defined mutant library. 0909IΔ*ethA* was constructed by unmarked gene deletion. Hemolysin-overexpressing *E. coli* strains were constructed by introducing into the wild-type strain (EIB202, CCTCC M208068) the plasmid of pSF4000-*hlyBACD* (Dhakal & Mulvey, 2012) into UPEC UT189, EHEC EDL933 (Burgos *et al*, 2009), EPEC (*E. coli* O26) (Schmidt *e*t al, 1995) and K12, respectively.

#### Hemolytic activity assay

Hemolytic activity was quantified using a microtiter assay developed previously(Aldick et al, 2007) with slight modifications. Indicated bacterial strains were cultured for 18–20 h and then pelleted, resuspended in PBS, and standardized at 600 nm to an OD of approximately 1.0. Next, 100 μL bacterial suspensions or OMV preparations were mixed with equal volumes of sheep erythrocytes in a 96-well plate. The plate was incubated at 37°C with gentle shaking for the indicated time periods and erythrocytes were pelleted by centrifugation (400 × *g*, 5 min). Clear supernatants were transferred to a fresh plate and absorbance at 540 nm was measured using a microplate reader (Dynex Technologies, Chantilly, VA, USA). Suspension buffer containing 0.1% Triton X-100 was served as a total lysis control. The percentage of hemolysis was calculated as follows: % hemolysis = (OD_540_ of samples - OD_540_ of background)/ (OD_540_ of total - OD_540_ of background) × 100.

#### Quantitative RT-PCR

RNA was extracted using an RNA isolation kit (Tiangen, Beijing, China). One microgram of each RNA sample was used for cDNA synthesis with the FastKing One Step RT-PCR Kit (Tiangen) and quantitative real-time PCR (RT-qPCR) was performed on an FTC-200 detector (Funglyn Biotech, Shanghai, China) using SuperReal PreMix Plus (SYBR Green) (Tiangen). The gene expression of bacterial *ethA* was evaluated for three biological replicates, and the data for each sample were expressed relative to the expression level of the 16S gene by using the 2^-△△CT^ method.

#### *Caspase-4^−/−^* cell line establishment

sgRNA (Oligo1: GACCGGGTCATCTCTGGCGTACTCC; Oligo2: AAACGGAGTACGCCAGAGATGACCC) was designed (http://zifit.partners.org) and cloned into LentiCRISPR v2 (AddGene, Cambridge, MA, USA; 52961), containing puromycin resistance. Lentiviral particles were prepared in HEK239T cells as previously described(van Zwicht et al,2016). HT-29 or Caco-2 cells were prepared in approximately 30% density and infected with lentiviral particles containing 10 µg/mL polybrene for 24 h. Transfected cells were selected with puromycin (2 µg/mL). Cells were diluted to 5 cells/mL and seeded to 96-well plates, followed by an expansion period to establish a new clonal cell line. The caspase-4^−/−^ cell lines were confirmed by western blotting using anti-caspase-4 antibody.

#### Cell culture and infections

Wild-type or *Caspase-4^−/−^* Caco-2, HT-29 and HeLa cells were seeded and grown to a density of ∼4×10^5^ cells per well in 12-well plates or ∼10^5^ cells per well in 24-well plates. For bacterial infection, cells were incubated with indicated *E. tarda* or *E. coli* strains at a MOI of 25 or 50, respectively. Infection was initiated by centrifuging the plate at 600 ×g for 10 min. After 50-min incubation at 35°C for *E. tarda* strains or 37°C for *E. coli* strains, the plates were washed once with PBS and then transferred into fresh medium containing 100 µg/mL gentamicin (Gm) to kill extracellular bacteria. The time point after antibiotic treatment was recorded as 0 h after infection. For OMV infection, cells were seeded and grown to a density of ∼10^5^ cells per well in 24-well plates, and then incubated with purified OMVs at 100 µg/10^5^ cells. Cell supernatants were harvested at the indicated time points for subsequent assays.

#### Cytokine and LDH release measurement

Aliquots of cellular supernatants were transferred into 96-well plates (round bottom) and centrifuged at 1000 ×g for 5 min. The supernatants were transferred to another 96-well plate (flat bottom), and the plate was subjected to the cytotoxicity assay using a CytoTox 96 assay kit (G1780, Promega, Madison, WI, USA) or an ELISA kit (eBioscience, San Diego, CA, USA) according to the manufacturer’s protocol. Each sample was tested in triplicate. Cytotoxicity was normalized to Triton X-100 treatment (100% of control), and LDH release from uninfected/untreated cells was used for background subtraction.

#### OMV isolation

OMVs were purified from *E. tarda* strains as described previously with minor modifications (Bielaszewska *et al*, 2014). Briefly, the bacterial strains were grown in 3000 mL of DMEM till OD_600_ of ∼1.5 and the bacteria-free supernatant was collected by centrifugation at 5000 ×*g* for 10 min at 4°C. This supernatant was further filtered through a 0.45 µm filter and concentrated using a spin concentrator (GE Healthcare, Little Chalfont, UK; molecular weight cut off = 30 kDa). Subsequently, OMVs were pelleted by ultracentrifugation at 284100 ×*g* for 1.5 h at 4°C in a Beckman NVTTM65 rotor (Brea, CA, USA). Isolated OMVs were resuspended in PBS, transferred to the bottom of a 13-mL ultracentrifugation tube (Beckman Coulter) and adjusted to 45% OptiPrep (Sigma-Aldrich, St. Louis, MO, USA) in a final volume of 2 mL. Different OptiPrep/PBS layers were sequentially added as follows: 2 mL of 40%, 2 mL of 35%, 2 mL of 30%, 2 mL of 25% and 1 mL of 20%. Gradients were centrifuged (284000 ×*g*, 16 h, 4°C) in a Beckman NVTTM65 rotor and the fractions of equal volumes (1 mL) were removed sequentially from the top. After removing the OptiPrep, OMVs were resuspended in 1000 µL sterile DMEM without phenol red. The protein content of purified OMVs was assessed by modified Bradford protein assay kit (Shenggong Biotech, Shanghai, China). OMV-free supernatants or OMVs, treated with proteinase K (Sigma) (5 µg/mL, 5 min) or not were separated by SDS-PAGE and immunoblotted with antibodies against OmpA, EthA, or RNAP.

#### OMV uptake assay

HeLa cells were seeded and grown to a density of ∼7 × 10^4^ cells per well in 24-well plates. Cells were pretreated with dyn (80 μM) or CD (1 μg/mL) for 30 min at 37°C and incubated with OMVs at 28 µg/10^5^ cells for 4 h. Cells were then washed, fixed, and quenched, and permeabilized/blocked with PBS containing 1 mg/mL saponin 10% and goat serum. OMVs were stained with anti-OmpA antibody and Alexa Fluor 488-conjugated goat anti-rabbit IgG. Actin was counterstained with TRITC phalloidin (Yeasen Biotech, Shanghai, China) and nuclei with DAPI (Beyotime, Jiangsu, China). Fixed samples were observed under a confocal microscope (Nikon, Tokyo, Japan; A1R).

#### Galectin assay

The eukaryotic expression plasmid for GFP tagged galectin-3 was constructed and introduced into HeLa cells by lentivirus transfection (Bielaszewska *et al*, 2014). Briefly, HeLa cells were seeded in 24-well plates at a density of 5 × 10^4^ cells per well in antibiotic-free medium and transfected with lentiviral particles for 24 h. Next, positively-transfected cells were pooled via treating cells with 1.5 µg/ mL of puromycin. The pooled cells were seeded and grown to a density of ∼10^5^ cells per well in 24-well plates. For bacterial infection, transfected cells were preincubated with EIPA (30 μM) or Dyn (80 µM) for 1 h before infection (DMSO pre-treatment as a control) and then challenged with different *E*. *tarda* strains at an MOI of 50. After 2.5 h-incubation at 35°C, the specks of galectin recruitment were observed under a confocal microscope (Nikon, A1R). For OMV infection, cells were incubated with purified OMVs at 100 µg/10^5^ cells. After 16-h incubation, the cells were washed twice with sterile PBS and fixed with 4% paraformaldehyde (PFA) at 25°C for 2 h, then washed in PBS and permeabilized with Triton X-100 (0.1% in PBS, 10 min at 25°C), and blocked in 5% bovine serum albumin. After overnight incubation with OmpA antibody, the cells were incubated with secondary fluorescent antibody for 1 h and DAPI was used for nuclear counterstaining. Fixed samples were observed under a confocal microscope (Nikon, A1R)

#### Cytosol extraction and LPS quantification

Subcellular cell fractions were extracted by a digitonin-based fractionation with modifications (Ramsby & Makowski, 2011). Briefly, *Caspase-4^−/−^* HeLa or Caco2 cells post incubation with indicated strain or purified OMVs were washed with sterile cold PBS 6 times on a platform shaker on ice to remove attached bacteria or OMVs. Subsequently, 250 or 100 µL of 0.005% digitonin extraction buffer was added to the bacteria-incubated cells in 12-well plates or OMV-incubated cells in 24-well plates, respectively. After 8 min, the supernatants were collected by centrifugation as the fraction containing cytosol, and the residuals were resuspended in 250 or 100 µL of 0.1% CHAPS buffer, respectively, as the fraction containing cell membrane, organelles and nucleus. Cytosol and residual fractions were subjected to the Limulus Amebocyte Lysate (LAL) assay (Associates of Cape Cod, East Falmouth, MA, USA) according to the manufacturer’s instructions to quantify LPS.

#### Mice and infections

C57BL/6J wild-type and *Caspase-11^−/−^* mice from Jackson Lab (6–8 weeks old) were bred under specific pathogen-free conditions. For oral infections, water and food were withdrawn 4 h before per os (p.o.) treatment with 20 mg/100 µL streptomycin per mouse. Afterward, animals were supplied with water and food ad libitum. At 20 h after streptomycin treatment, water and food were withdrawn again for 4 h before the mice were orally infected with 5 × 10^7^ CFU/g of EIB202 or 0909I suspension in 200 µL PBS, or treated with sterile PBS (control). Thereafter, drinking water ad libitum was offered immediately and food 2 h post-infection. At the indicated times points, mice were sacrificed and the tissue samples from the intestinal tracts, kidneys, spleens, and livers were removed for analysis.

#### Bacterial counts and histopathology analysis

Collected tissues or organs were homogenized in PBS (pH 7.4) and the dilutions were plated on Deoxycholate Hydrogen Sulfide Lactose (DHL) agar plates for CFU counting. The cecum and colon were collected in 10% neutral-buffered formalin for histological analyses. Tissue pathology was blindly scored by two researchers using hematoxylin and eosin-stained sections (6 µm). The scoring criteria for submucosal edema, PMN infiltration into the lamina propria, gobletcell loss and epithelial integrity was conducted as previously described by Barthel *et al*. (Barthel *et al*, 2014). In addition, inflammatory focal infiltration (IFI) within cross-section was scored at five levels (0, none; 1, inflammation occurrence, but without apparent IFI; 2, with apparent IFI; 3, more than one to three IFIs; 4, with more than three IFIs). The cumulative scoring range was 0–17.

#### Immunohistochemistry of gut sections

Cryosections (8 µm) of OCT (Leica)-embedded tissue samples from the cecum and colon of each animal were mounted on glass slides (Thermo Scientific, Waltham, MA, USA), fixed with 4% paraformaldehyde (PFA) at at 25°C for 2 h, washed in PBS, and blocked with blocked in 5% bovine serum albumin for 15 min. Next, the cryosections were incubated overnight with monoclonal rat anti-GSDMDC1 (1:50). Secondary fluorescent antibodies were added for 1 h, washed with PBS, TRITC phalloidin (Yeasen, Shanghai, China) was added for 30 min to counterstain actin, and DAPI was used for nuclear counterstaining. Fixed samples were observed under a confocal microscope (Nikon, A1R).

#### Mucosal and serum cytokine measurements

Cecal and colonic tissues were removed from mice 24 h after infection with *E. tarda* strains. Tissues were washed free of luminal contents and then incubated in DMEM (supplemented with penicillin and streptomycin at 1% and streptomycin at 1 mg/mL) for 6 h, and the supernatants, along with the serum samples, were collected for IL-18 detection by ELISA following the manufacturer’s protocols.

#### Antibodies

Rat anti-GSDMDC1 (1:50; sc-393656; Santa Cruz Biotechnology, Dallas, TX, USA) was used for immunohistochemistry of mice gut sections. Rabbit anti-*E. tarda* OmpA polyclonal antibody (1:500; custom-made; Genscript Biotech, Piscataway, NJ, USA), rabbit anti-*E. tarda* EthA polyclonal antibody (1: 1000; custom-made; Genscript), and mouse anti-RNAP monoclonal antibody (1:5000; Santa Cruz Biotechnology) were used for western blotting or immunofluorescence.

